# The mutational decay of male and hermaphrodite competitive fitness in the androdioecious nematode *C. elegans*, in which males are naturally rare

**DOI:** 10.1101/115568

**Authors:** Shu-Dan Yeh, Ayush Saxena, Timothy Crombie, Dorian Feistel, Lindsay M. Johnson, Isabel Lam, Jennifer Lam, Sayran Saber, Charles F. Baer

## Abstract

Androdioecious Caenorhabditis have a high frequency of self-compatible hermaphrodites and a low frequency of males. The effects of mutations on male fitness are of interest for two reasons. First, when males are rare, selection on male-specific mutations is less efficient than in hermaphrodites. Second, males may present a larger mutational target than hermaphrodites because of the different ways in which fitness accrues in the two sexes.

We report the first estimates of male-specific mutational effects in an androdioecious organism. The rate of male-specific inviable or sterile mutations is ≤ 5 x 10^−4^/generation, below the rate at which males would be lost solely due to those kinds of mutations. The rate of mutational decay of male competitive fitness is ~0.17%/generation; that of hermaphrodite competitive fitness is ~0.11%/generation. The point estimate of ~1.5X faster rate of mutational decay of male fitness is nearly identical to the same ratio in Drosophila. Estimates of mutational variance (VM) for male mating success and competitive fitness are not significantly different from zero, whereas VM for hermaphrodite competitive fitness is similar to that of non-competitive fitness. The discrepancy between the two sexes is probably due to the greater inherent variability of mating relative to internal self-fertilization.

## Introduction

Several species of nematodes in the genus Caenorhabditis, among them the well-known *C. elegans,* have evolved an androdioecious mating system in which self-fertilizing hermaphrodites are very common and males are very rare. In *C. elegans,* for example, the frequency of outcrossing (= male-female mating, because hermaphrodites cannot mate with each other) is thought to be on the order of 1% or less, perhaps much less [1]. Dioecy is ancestral in the genus and most species in the genus are dioecious, although androdioecy has evolved independently at least three times [2]. Moreover, at least in *C. elegans,* androdioecy appears to have evolved quite recently [3].

Sex determination in Caenorhabditis is an XO type chromosomal system, with females/hermaphrodites having two copies of the X chromosome and males having a single X chromosome [4]. Laboratory populations of *C. elegans* kept under constant conditions in which the frequency of males is initially elevated above the background consistently lose males, until the frequency of males equilibrates at the frequency of non-disjunction of the X [5]. The frequency of males varies among strains [6] and depends on environmental conditions, averaging about 0.1% under standard husbandry conditions in the N2 strain. However, laboratory populations exposed to variable selection can keep males at significantly higher frequencies [7], consistent with the idea that recombination facilitates adaptive evolution.

In an androdioecioius population, selection on male function is (*i*) necessarily weaker than selection on hermaphrodite function and (*ii*) weaker than selection on male or female function in a dioecious population, because in the absence of males (or females) a dioecious population immediately goes extinct whereas an androdioecious population plods on even in the complete absence of males. Moreover, although selection typically favors an equal sex-ratio in a randomly mating dioecious population [8], that is not generally true in a partially selfing population [9].

Importantly, the rarer males become, the less efficient is selection against genes with male-specific effects on fitness. For example, if males represent 0.1% of the population, as in a typical *C. elegans* lab population, an autosomal gene with effects only on male fitness will find itself in a male - and thus under selection - 0.1% of the time and in a hermaphrodite - and thus free from selection - 99.9% of the time. If the selection coefficient on that gene is 10% in males and 0 in females, the average selection coefficient at the gene will be 0.01%. The effective population size, *N_e_*, of *C. elegans* is thought to be on the order of 10,000 [10], so an allele with those sex-specific selection coefficients will have an average selection coefficient of approximately 1/*N_e_*, roughly the boundary of effective neutrality [11]. Thus, the rarer males become, the stronger selection must be on male function to overcome random genetic drift.

These features of selection in androdioecious populations lead to a chicken-and-egg question with respect to the rarity of males in androdioecious Caenorhabditis: are males rare because the sex-ratio is near an evolutionary optimum, or are males on their way out, doomed to ultimately succumb to the ravages of deleterious mutation? Or perhaps both. A quantitative answer to that question requires an estimate of the distribution of effects of mutations affecting male fitness, both with respect to mutations that render males non-viable or sterile, and the effects on the ability of males to mate and for male sperm to compete with hermaphrodite sperm. Several elements of male fitness have been shown to be genetically variable among wild isolates of *C. elegans* [12].

Unfortunately, the distribution of fitness effects (DFE) on individual traits is very difficult to quantify reliably [13]. More tractable measures of the vulnerability of a trait to the cumulative effects of mutation are (*i*) the rate of change of the trait mean due to the accumulation of spontaneous mutations (the “mutational bias”, ΔM) and (*ii*) the rate of increase in genetic variance (the “mutational variance”, VM). These quantities can be used to quantify the relative mutability of traits and populations.

Here we report on an experiment designed to estimate the cumulative effects of spontaneous mutations on male-male competitive fitness in a set of *C. elegans* mutation accumulation (MA) lines which were propagated by single hermaphrodite descent for approximately 250 generations. In this context, male function constitutes a truly neutral trait, since chromosomes were (almost) never passed through males. Cumulative effects of mutations on many hermaphrodite traits have been previously reported for these and other *C. elegans* MA lines (summarized in [14]), providing a robust baseline against which to compare male mutational properties. In addition, we report new results on the cumulative mutational effects on hermaphrodite-hermaphrodite competitive fitness.

The results will shed light on two questions of interest. First, the frequency of MA lines for which fertile males can be obtained provides a rough upper bound on the class of mutations resulting in “zero male fitness”; this class comprises male-specific lethal and male-sterile mutations. There are surprisingly few published estimates of the rate of mutation to alleles of zero male fitness. Mukai et al. [15] reported the frequency of matings of *Drosophila melanogaster* MA lines resulting in sterility was “below 2%”. Willis [16] reported that a large fraction of inbreeding depression in *Mimulus gutatus* (~30%) could be attributed to male-sterile mutations, but actual rates could not be quantified.

Second, male components of fitness may be particularly vulnerable to the effects of deleterious mutations, for two reasons. Male mating success is much closer to a winner-take-all game than other components of fitness such as female fecundity or egg-to-adult viability (unless predation is an important cause of mortality). Small differences in performance may be magnified into a win-lose outcome, with the winner mating and the loser not mating. Also, the “genic capture” hypothesis [17] predicts that sexual selection acts at least indirectly on male condition, in which case the mutational target of male fitness is potentially very large. Given the infrequency of outcrossing in *C. elegans,* it is arguable whether sexual selection is important to the evolution of the species. However, given that (*i*) outcrossing is the ancestral state in the genus, (*ii*) androdioecy seems to have evolved relatively recently in *C. elegans,* and (*iii*) the basic biology of mating and fertilization appears similar throughout the genus, it seems reasonable that the cumulative mutational effects on male fitness in *C. elegans* would at least approximately reflect those in dioecious species.

To date, the only published estimates of cumulative mutational effects on male fitness in any animal come from *Drosophila melanogaster* [18-20], in which effects on male fitness are generally somewhat greater than those on female fitness.

## Methods and Materials

### Mutation accumulation (MA)

The details of the construction and maintenance of the MA lines have been reported elsewhere [21]. Briefly, 100 replicate lines were initiated from a highly homozygous population of the N2 strain of *C. elegans* and maintained by serial transfer of a single immature hermaphrodite every generation for approximately 250 generations, at which point each MA line was cryopreserved. The common ancestor (G0) of each set of MA lines was cryopreserved at the outset of the experiment.

### Recovery of males from MA lines

Beginning in the winter of 2015, cryopreserved 250- generation (G250) MA lines were thawed and replicate populations collected on standard 60 mm NGM agar plates. For lines in which males were not present on the thawed plate, we attempted to generate males using a standard heat shock protocol to induce non-disjunction of the X [22]. If males were obtained but pairings with hermaphrodites failed to produce male progeny, after three heat shock attempts the MA line was characterized as producing sterile males. If no males were obtained after three heat shock attempts, the MA line was characterized as incapable of producing males. Once males were obtained, a single male was paired with three young L4-stage hermaphrodites on a 35 mm NGM agar plate seeded with OP50 strain *E. coli,* and the progeny split into two 100 mm NGM agar plates seeded with OP50, grown until food was just exhausted, and cryopreserved. A set of 46 male-segregating G0 “pseudolines” was constructed in the same way and cryopreserved at the same time.

### Male competitive fitness assay

Male-male competitive fitness was assayed by pairing a focal male (G0 ancestor or MA) with a marked competitor male of the ST-2 strain (homozygous for the dominant ncls2 pH20::GFP reporter allele on an N2 genetic background) and a male-sterile hermaphrodite homozygous for a recessive null allele at the *fog-2* locus [*fog-2*(*q71*)]*. fog-2* is a recessive mutation that destroys spermatogenesis in hermaphrodites, thereby rendering hermaphrodites functionally female [23]. To minimize segregating variance in the maternal stock, we backcrossed the *fog-2*(*q71*) mutant allele into the ancestral N2 genetic background for ten generations prior to initiating the competitor population from a cross of a ST-2 male with a *fog-2* female.

The assay was performed in two blocks, in the same conditions as the MA phase of the experiment (plates seeded with 100 μl of an overnight culture of the OP50 strain of *E. coli* as food, incubated at 20°), with the exception that the assay plates were 40% agarose (NGMA) rather than 100% agar, to prevent worms from burying in the substrate. Each MA line was included in each block. Assay blocks were initiated by thawing MA lines and G0 pseudolines, followed by one generation of male recovery in ten replicate 35 mm plates. Each replicate plate contained two or three males and three young hermaphrodites. All lines were thawed and replicated on the same day. Replicate plates were assigned random numbers and were subsequently handled in order by random number. On the third day after the replicate plates were initiated, competition assay plates were initiated by transfer of a single young focal male from each replicate plate and a single similarly-staged competitor male from a stock plate. The two males were allowed to acclimate to the plate for one day, at which time a female was introduced to the plate at a location approximately intermediate between the two males. Two days after the introduction of the female, the three adult worms were removed from the plate and their progeny allowed to grow for another two days. In Block 1, after the two-day growth period, plates were stored at 4° C for four days prior to counting. In Block 2, worms were counted directly after the two-day growth period without refrigeration.

Worms were counted with the aid of a Union Biometrica BioSorter™ large-particle flow cytometer (aka, a “worm sorter”) equipped with the LP Sampler™ microtiter plate sampler. The detailed counting protocol is given in Supplementary Appendix A1. Worms were washed from the competition plates in approximately 1.5 ml of M9 buffer into 1.5 ml microcentrifuge tubes. Tubes were centrifuged at ~2K x *g* for 1 minute, the supernatant decanted, and the pelleted worms resuspended in 100 μl of M9 and pipetted into a well in a 96-well microtiter plate, which was counted with the BioSorter as outlined in Supplementary Appendix A1.

### Hermaphrodite competitive fitness assay

Competitive fitness of hermaphrodites was assayed in two blocks beginning in May, 2005. At the outset of each block, the cryopreserved G0 ancestor of the MA lines was thawed and 20 replicate populations initiated from a single L3/L4 stage worm placed on a standard 60 mm NGM agar plate seeded with 100 μl of an overnight culture of the OP50 strain of *E. coli.* These populations are referred to as “pseudolines” and designated the P0 generation. Seven L3/L4 stage offspring from each pseudoline were transferred singly to new plates, designated the P1 generation. From this point on, pseudolines were treated identically to the MA lines. G250 MA lines were thawed and seven revived L3/L4 stage worms from each line were placed individually on standard 60 mm NGM plates, labeled P1. All P1 plates were assigned a unique random number and all subsequent experimental manipulations were performed in sequence by random number. All replicate populations were maintained for two more generations (P2-P3) by transfer of a single L3/L4 stage offspring at four-day intervals to control for parental and grandparental effects. At the same time, we thawed a replicate of the GFP-marked competitor strain ST-2 and made several large replicate populations by transferring a chunk from the initial plate to a new 100 mm plate.

On the second day after the P3 worm began reproduction, a competition plate for each replicate was set up by transferring a single L1-stage larva from the P3 plate and a single L1-stage ST-2 competitor onto a 60 mm NGM agar plate seeded with 100 μl of the HB101 strain of *E. coli* and supplemented with nystatin to retard fungal contamination. Competition plates were incubated at 20° C for eight days, at which point food was exhausted. Worms were washed from competition plates in cold M9 buffer, settled on ice and 100 μl of the settled worms transferred into a drop of glycerol on the lid of an empty 60 mm agar dish and the bottom of the empty dish pressed into the lid. The glycerol immobilizes the worms and pressing them between halves of the plate puts them into the same focal plane. We took two pictures of each plate at 40X magnification through a Leica MZ75 dissecting microscope fitted with a 100 W mercury arc lamp and epifluorescence GFP filter cube (470/40 nm excitation filter, 525/50 nm emission filter) using a Leica DFC280 camera connected to a computer running the Leica IM50 software (Leica Microsystems Imaging Solutions Ltd). The first picture used the arc lamp and GFP filter cube (called the “green” image) and the second, taken immediately afterwards, used transmitted white light (called the “white” image). All worms are visible in the white image, whereas wild-type (non-GFP) worms appear only very faintly in the green image (Supplementary Figure S1). The difference between the number of worms in a white image and in the matching green image is the number of focal worms in the sample.

Images were imported into ImageJ software (http://rsb.info.nih.gov/ij/) and worms were counted as follows. If there appeared to be fewer than 200 worms visible in a white image, we first counted every worm in the white image and then each worm visible in the accompanying green image. If there appeared to be > 200 worms in the white image we drew a rectangle around approximately 200 worms and counted them. We then pasted the same rectangle in the green image and counted the worms visible within the rectangle.

### Data Analysis

#### i) Measures of competitive fitness

Competitive fitness has two components: (1) did the focal individual reproduce at all? If not, relative fitness is zero regardless of the number of offspring of the competitor, and (2) given that the focal individual did reproduce, what fraction of the offspring belong to the focal individual? These considerations apply both to male-male competitive fitness and to hermaphrodite-hermaphrodite competitive fitness. Given that a focal individual did reproduce, the ratio *p/(1-p)* is related to competitive fitness by the relationship

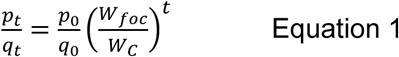

[24, equation 17.2], where *t* represents the number of generations in the fitness assay (NOT the number of MA generations), *p*_0_ is the frequency of the focal type (G0 or control) at the beginning of the assay, *p_t_* is the frequency of the focal type (G0 control or MA) at the conclusion of the assay, *q* = 1-*p*, *W_foc_* is the absolute fitness of the focal type, and *W_C_* is the absolute fitness of the competitor. Each trial was started with one focal worm and one competitor, so the ratio 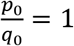. We refer to the ratio *p*/(1-*p*) as the “competitive index”, *CI* [25]. *CI* provides a measure of fitness of the focal type relative to the competitor, raised to the power *t*. All analyses of *CI* were performed on natural log-transformed data.

#### ii) Probability of reproduction, *П*

Probability of reproduction is a binary trait. If a focal worm reproduced the replicate is scored as a success (“event=1”); if the focal worm did not reproduce it is scored as a failure (“event=0”). Data were analyzed by Generalized Linear Mixed Model (GLMM) with estimation by Residual Subject-specific Pseudolikelihood (RSPL) as implemented in the GLIMMIX procedure of SAS v.9.4 with a logit link function and a random residual. Treatment (MA vs. Control) is a fixed effect and Line and Replicate (nested within Line) are random effects. Block is a random effect in principle. However, pseudolikelihoods are not appropriate criteria for model selection (e.g., by AIC; [26]), so rather than include or exclude variance components including block on the basis of estimates for which there is little power (because *n*=2), we chose to model block as a fixed effect for this analysis. It is common in the analysis of MA fitness assays to treat block as a fixed effect when the number of blocks is small (e.g., [27, 28]).

Each line (MA and G0 pseudoline) was assayed for male-male competitive fitness, *П_M_*, in each of the two assay blocks. The full model is written as:

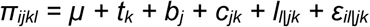

where *π_ijkl_* is a binary variate scored as 1 if the focal worm produced at least two offspring and 0 if it did not, *μ* is the overall mean, *t_k_* is the fixed effect of treatment *k* (G0 or MA), *b_j_* is the fixed effect of block *j*, *C_jk_* is the fixed effect of the treatment by block interaction, *I_l_*_\*jk*_ is the random effect of line (or pseudoline) *l*, conditioned on block and treatment, and *ε_il_*_\*jk*_ is the random residual, conditioned on block and treatment. Random effects were estimated separately for each block/treatment combination by means of the GROUP option in the RANDOM statement of the GLIMMIX procedure [26]. Significance of fixed effects was determined by F-test of Type III sums of squares, with degrees of freedom determined by the Kenward-Roger method [29].

Hermaphrodite probability of reproduction, *π_H_*, was modelled similarly, with the exception that each line (or pseudoline) was represented in only one of the two assay blocks, so line is nested within block. The distribution of *π_H_* was strongly left-skewed, so means and standard errors were calculated by an empirical bootstrap procedure [30, 31]. Resampled datasets were constructed by resampling lines within blocks, followed by estimation of means and variance components from the GLMM described above.

#### Competitive Index (*CI*)

*(i) Males.* Given that both male worms – the focal worm and the competitor – sired at least 2% of the offspring on the competition plate, we analyzed false-positive corrected male-male *CI* (*CI_M_*) using a standard general linear model (GLM) as implemented in the MIXED procedure of SAS v. 9.4. The correction for false positives is explained in Supplementary Appendix A1. Studentized residuals of natural log-transformed data were scrutinized for outliers by eye against a Q-Q plot. After removal of three outliers (*n =* 671), the data were initially fit to the linear model:

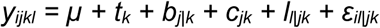

where *y_ijkl_* is the log(*CI*) of the individual replicate and the independent variables are defined as in the previous section. Block and Line-by-block interaction were modelled as random effects in this analysis. Variance components of random effects were estimated by restricted maximum likelihood (REML). The among-block component of variance was estimated separately for each treatment and the among-line and among-replicate (nested within line) components were estimated separately for each treatment/block combination by means of the GROUP option in the RANDOM or REPEATED statement of the MIXED procedure [32].

We first analyzed the full model above, then sequentially simplified the model by first pooling the random effects across grouping levels (e.g., estimating a single among-line variance rather than estimating it separately for each block) and then removing the effect entirely. The model with the smallest corrected AIC (AICc) was chosen as the best model, and significance of the fixed effect of treatment (MA or G0) in that model was determined by F-test of Type III sums of squares, with degrees of freedom determined by the Kenward-Roger method [29]. If two models had equal AICc, the simpler model was chosen as the best model. AICc’s of the models tested are given in Supplementary Table S1. In addition, we calculated empirical bootstrap estimates of the mean and standard error of *CI_M_*, resampling over lines within blocks followed by estimation of means and variance components from the GLM described above [30, 31].

Hermaphrodite *CI*, *CI_H_* was modelled analogously to *CI_M_*, with the exception that each line (or pseudoline) was represented in only one of the two assay blocks, therefore line is nested within block and there is no line-by-block interaction term. Outliers were identified as for males; five outliers (*n* = 727) were removed prior to further analysis.

#### Mutational Bias

The mutational bias is the per-generation rate of change in the trait mean. The slope of the regression of trait mean on generation of MA is often designated *R_M_* [33]; the per-generation change scaled as a fraction of the ancestral (G0) trait mean is often referred to as 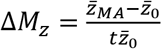, where *z̅_MA_* and *z̅*_0_ represent the MA and ancestral trait means and *t* is the number of generations of MA. For *П_M_* and *П_Η_*, MA and G0 means were estimated by least squares, given the general linear mixed model, and ΔM_П_ calculated directly from the least-squares means. *CI* is on a logarithmic scale so mean-standardization of the data is not appropriate because *CI* can be negative or positive. For *CI*, 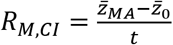 and the mutational bias represents the per-generation change in competitive fitness of the focal genotype relative to the competitor strain. *R_M_*,*CI* can be related to the per-generation mutational change in relative fitness *per se,* ΔM_*w*_, from Equation 1, as explained below in the Results.

#### Mutational Variance (*VM*)

The per-generation increase in genetic variance resulting from new mutations, *VM,* is equal to the product of the per-genome, per-generation mutation rate (*U*) and the square of the average effect of a mutation on the trait of interest, α, i.e., *VM=Ua*^2^ [34]. In this experiment, MA lines are assumed to be homozygous, in which case VM is equal to half the increase in the among-line component of phenotypic variance, divided by the number of generations of MA, i.e, 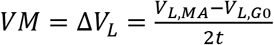, where *V_L,MA_* is the variance among MA lines, *V_L_*_,*G*0_ is the variance among the G0 pseudolines, and *t* is the number of generations of MA ([35], p. 330). For all traits (*π* and *CI*), VL is the among-line component of variance of the treatment group (MA or G0) extracted from the relevant GLM or GLMM.

VM is typically scaled either relative to the residual (environmental) variance, VE or to the square of the trait mean, *z̅.* The ratio VM/VE is the mutational heritability 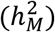, and the ratio 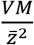 is sometimes called the mutational evolvability (*I_M_*) and is equivalent to the squared mutational coefficient of variation [36]. Usually, VM is scaled relative to the ancestral (G0) mean, but if the mean changes substantially over the course of MA (i.e., ΔM *≠* 0), it is more meaningful to scale each group (G0 and MA) by its own mean [21]. Scaling by the group means is nearly equivalent to calculating *ΔV_L_* from log-transformed data [37]. *CI* is on a logarithmic scale and cannot be mean-standardized. *ΔV_L_* for *CI_M_* and for *П_M_* and *П_Η_* is not significantly different from 0 (see Results), so scaling is irrelevant.

## Results

### Rate of mutations with Zero Male Fitness

Of the 60 of the original 100 MA lines remaining in 2014, we were able to obtain fertile males from 53. We assume that the seven lines for which we were unable to obtain fertile males carry at least one mutation that leads to Zero Male Fitness (ZMF, i.e. inviable or sterile), and that these mutations follow a Poisson distribution - analogous to lethal equivalents [38]. With those assumptions, the expected proportion of lines with no mutations (*p*_0_, i.e., the number of lines that produced fertile males) is: *p*_0_ = *e^-m^*, where *m* is the expected number of mutations carried by a line [39]. The expected number of mutations *m* = *μt*, where *μ* is the per-generation rate of ZMF mutations and *t* is the number of generations of MA. Thus, (53/60) = e^−250*μ*^, so *μ_ZMF_* ≈ 5×10^−4^/generation, about half the lower-bound estimate on the frequency of males. If the ZMF mutation rate is greater than the frequency of males, the average chromosome will take a male-sterilizing hit before the next time it winds up in a male, leading to the loss of males by “error catastrophe” [40]. The same calculation from data reported in [41] gives an estimate of *μ_ZMF_* ≈ 3×10^−4^/generation.

### Male-Male Competitive Fitness

*(i) Probability of mating* (*π_M_*). After 250 generations of completely relaxed selection, MA males are significantly less likely to successfully mate under competitive conditions than are their unmutated G0 ancestors (F_1,131.8_ = 12.51, P<0.0001). Averaged over the two blocks, the probability that a G0 male mated successfully (defined as an estimated frequency of offspring sired > 2%) was ~90% (*π̅_M_ =* 0.912 ± 0.019) whereas the probability that a MA male mated significantly declined to ~75% (*π̅_M_ =* 0.757 ± 0.023). Scaled relative to the G0 mean, the probability of a male successfully mating under the assay conditions decreased by about 0.06% per generation (ΔM_*П*_ = −0.622 ± 0.159 x 10^−3^/generation; Table 1; distributions of line means are shown in Supplementary Figure S2).

**Table 1.** Evolution of trait means, averaged across blocks; standard errors in parentheses. Sample sizes are given in Table 2. Values of *R_M_* significantly different from zero (P<0.05) shown in bold type. See Methods for abbreviations and details of calculations. ΔM of CI is calculated for back-transformed data. * - Assumes *t* = 2 generations of reproduction † - ΔM calculated for *CI*, not log(*CI*)

| | $\pi_M$ | $\pi_H$ | $\log(CI_M)$ | $\log(CI_H)$ |
| --- | --- | --- | --- | --- |
| G0 | 0.912 (0.016) | 0.987 (0.008) | -0.145 (0.074) | 0.862 (0.331) |
| MA | 0.757 (0.023) | 0.982 (0.005) | -0.461 (0.095) | 0.228 (0.338) |
| $R_M$ (x 10<sup>3</sup>/gen) | **-0.622** (0.159) | -0.0192 (0.050) | **-1.26** (0.48) | **-2.54** (0.49) |
| $\Delta M$ (x 10<sup>3</sup>/gen) | -0.682 | -0.0195 | -1.04<sup>†</sup> | -1.09<sup>*,†</sup> |

In neither block did VM of *π_M_* differ significantly from 0. In the first block, the RSPL estimate of VL among G0 pseudolines was greater than VL among MA lines; in the second block VL was greater in the MA lines but the difference was not significant (Table 2). *(ii) Competitive Index (CIM).* When the estimated frequency of offspring sired by the focal male was at least 2%, MA males sired a smaller fraction of offspring than did their unmutated G0 ancestors [log(*CI_M_*,_*G0*_) = −0.145 ± 0.074; log(*CI_M_*_,*MA*_) = −0.461 ± 0.095; standard errors represented by the standard deviation of the empirical bootstrap distribution]. The best-fit linear model includes a separate among-block component of variance for each treatment (Supplementary Table S1), and under that model the change in the trait mean is not significantly different from zero (F_*1,1.38*_ =1.67, P > 0.37). However, when the among-block variance is pooled over the two treatments, the change in the mean *CIM* becomes significant (F_*1,603*_ = 5.57, P < 0.02). The lack of significance in the best-fit model potentially represents Type II error resulting from having to estimate a variance component with *n*=2. To test that possibility, we estimated the change in the trait mean from the mean of 1000 bootstrap replicates, resulting in an empirical P < 0.007. Averaged over the two blocks, log(*CI_M_*) declined by slightly more than 0.1% per generation (*R_M,CI,M_* = −1.26 ± 0.48 x 10^−3^ /generation; Table 1; distributions of line means are shown in Supplementary Figure S3).

**Table 2.**
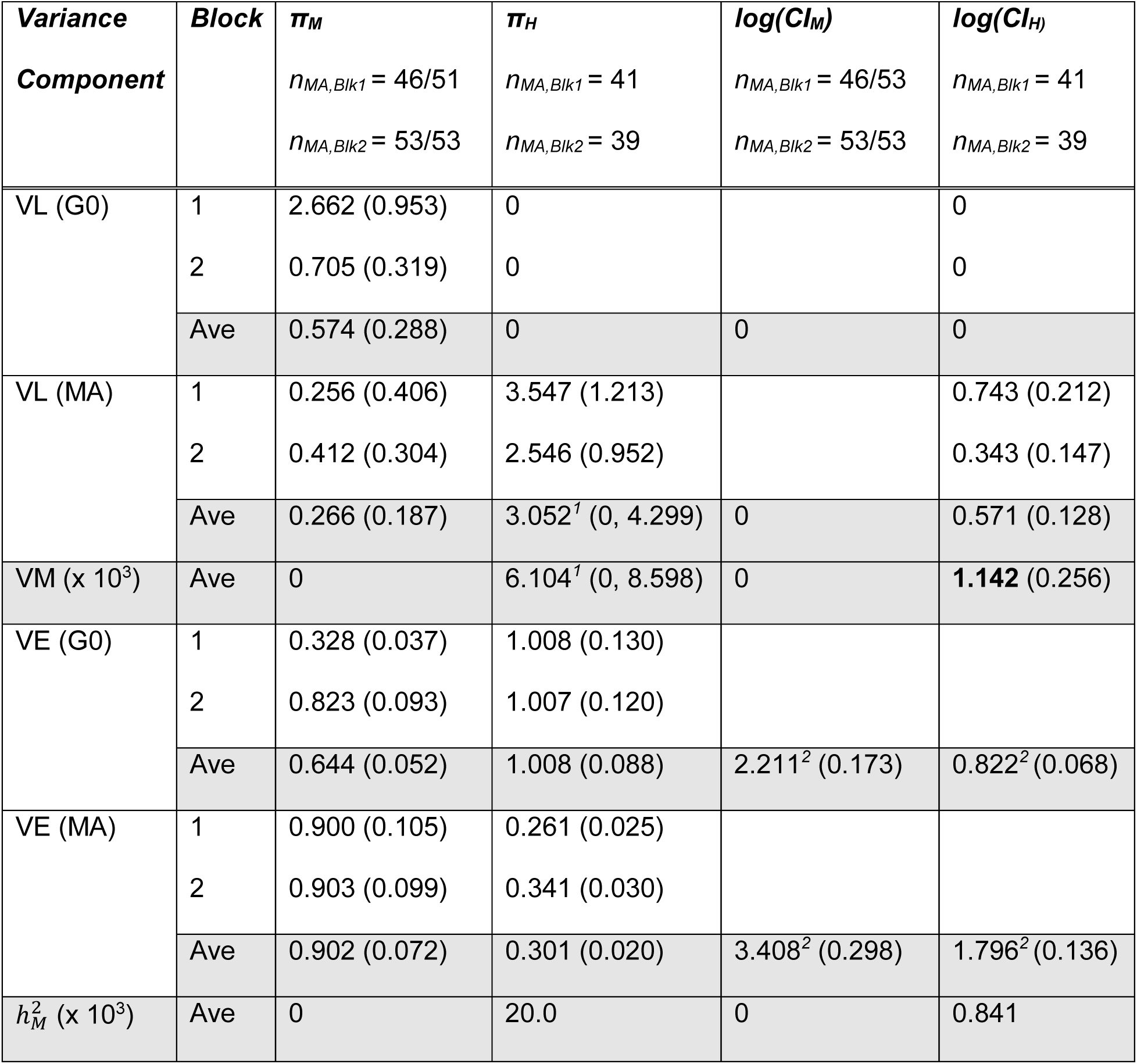
Variances. Standard errors in parentheses. See Methods for abbreviations and details of calculations. For male traits, n*_μA,Blk_*. is the fraction of the 53 MA lines that were included in that block. For hermaphrodite traits, each block had a different set of MA lines, out of 80 total lines. VM significantly greater than 0 (P<0.05) shown in bold type. ^1^ - 95% bootstrap confidence interval in parentheses ^2^ - Best-fit model includes residual variances for each treatment (G0, MA) pooled over blocks.

| **Variance Component** | **Block** | $\pi_M$<br>$n_{MA,Blk1} = 46/51$<br>$n_{MA,Blk2} = 53/53$ | $\pi_H$<br>$n_{MA,Blk1} = 41$<br>$n_{MA,Blk2} = 39$ | $\log(CI_M)$<br>$n_{MA,Blk1} = 46/53$<br>$n_{MA,Blk2} = 53/53$ | $\log(CI_H)$<br>$n_{MA,Blk1} = 41$<br>$n_{MA,Blk2} = 39$ |
| --- | --- | --- | --- | --- | --- |
| VL (G0) | 1 | 2.662 (0.953) | 0 |  | 0 |
|  | 2 | 0.705 (0.319) | 0 |  | 0 |
|  | Ave | 0.574 (0.288) | 0 | 0 | 0 |
| VL (MA) | 1 | 0.256 (0.406) | 3.547 (1.213) |  | 0.743 (0.212) |
|  | 2 | 0.412 (0.304) | 2.546 (0.952) |  | 0.343 (0.147) |
|  | Ave | 0.266 (0.187) | 3.052<sup>1</sup> (0, 4.299) | 0 | 0.571 (0.128) |
| VM (x 10<sup>3</sup>) | Ave | 0 | 6.104<sup>1</sup> (0, 8.598) | 0 | **1.142** (0.256) |
| VE (G0) | 1 | 0.328 (0.037) | 1.008 (0.130) |  |  |
|  | 2 | 0.823 (0.093) | 1.007 (0.120) |  |  |
|  | Ave | 0.644 (0.052) | 1.008 (0.088) | 2.211<sup>2</sup> (0.173) | 0.822<sup>2</sup> (0.068) |
| VE (MA) | 1 | 0.900 (0.105) | 0.261 (0.025) |  |  |
|  | 2 | 0.903 (0.099) | 0.341 (0.030) |  |  |
|  | Ave | 0.902 (0.072) | 0.301 (0.020) | 3.408<sup>2</sup> (0.298) | 1.796<sup>2</sup> (0.136) |
| $h_M^2$ (x 10<sup>3</sup>) | Ave | 0 | 20.0 | 0 | 0.841 |

As noted, *CI* cannot be directly mean-standardized. However, *CI* is related to relative fitness by equation [1] above. ΔM*_W_* can be calculated from 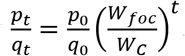, where 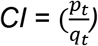, p_0_ = q_0_ = 0.5, and *t* = 1, thus 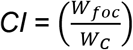. The ratio of the fitnesses relative to the competitor (designated by a capital *W*), 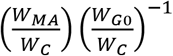 gives the fitness of the MA lines relative to that of the G0 ancestor (designated by a lower-case *w*), 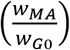. The ratio 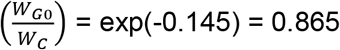 and 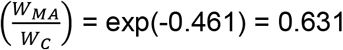, so 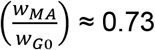. Thus, male competitive fitness relative to the ST-2 competitor declined by about 27% over the course of 250 generations of MA, or by about - 1.04 x 10^−3^/generation when scaled as a fraction of the G0 mean.

The REML estimate of VL for log(*CI_M_*) for both G0 pseudolines and MA lines is zero in each block (Table 2), and the distributions of line means are similar in the two groups (Supplementary Figure S3). Taken at face value, a change in the mean coupled with no change in the among-line variance implies that each line changed at the same rate, or at least at rates that were indistinguishable. A more plausible explanation is that the true genetic variance is small relative to the environmental variance (which includes experimental error) and the sample sizes employed here were not large enough to provide power to detect small differences. In the male fitness assay, each line was initially replicated tenfold, five replicates per block. In the hermaphrodite competitive fitness assay, in which ΔVL for log(*CI_H_*) is highly significant (P<0.0001; see next section), each line was replicated sevenfold, but in only one of the two blocks.

### Hermaphrodite-Hermaphrodite competitive fitness

#### (i) Probability of reproducing

(*П_H_*). The probability of a hermaphrodite reproducing was high (>98% for both G0 and MA), and changed little over 250 generations of MA (ΔM = −1.9 × 10^−5^/generation; T able 1; distributions of line means are shown in Supplementary Figure S2). The mutational heritability is large (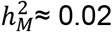; Table 2), but the distribution of *П_Η_* among lines is highly left-skewed (median *П_Η_* = 1; Supplementary Figure S2) so the point estimate of *h*^2^_*M*_ is probably not very meaningful.

It is likely that some GFP-marked competitors were misidentified as focal types (non-GFP) in our image analysis, which could potentially inflate the apparent probability of reproduction. However, these values of *π_H_* are nearly identical to the probability of reproduction of hermaphrodites in a different experiment in which hermaphrodites of the same set of MA lines were allowed to reproduce in non-competitive conditions (*π_H_* > 97% for both G0 and MA; ΔM = - 3.3 x 10^−5^/generation; reanalysis of data in [30]). Thus, the very high rate of reproduction does not appear to be an experimental artifact.

#### (ii) *Competitive Index* (*CI_H_*)

Mean *CI_H_* declined significantly over the course of 250 generations of MA [log(*CI_H_*_,*G0*_) = 0.862 ± 0.331; *log*(*CI_H,MA_*) = 0.228 ± 0.338; F_1,169_ = 26.97, P<0.0001; Table 1; distributions of line means are shown in Supplementary Figure S3]. *R_M,CI,H_* calculated from the slope of the regression of log(*CI_H_*) on generation of MA is −2.54 ± 0.49 x 10^−3^ per-generation.

From equation [1], 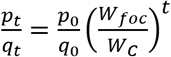, where *CI =* 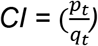, *p*_0_ = *q*_0_ = 0.5 and here *t* is equal to the number of generations of reproduction the population underwent over the course of the eight-day assay. Therefore, 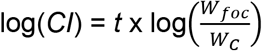, so 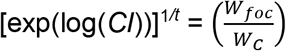, and the ratio of the fitnesses relative to the competitor, 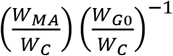 gives the fitness of the MA lines relative to that of the G0 ancestor, 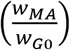.

We cannot be certain about the exact number of generations of reproduction, except that three is the upper bound (based on the worm’s life cycle), and the true number is probably close to two. The basis for that judgment is this: if the average worm produces 200 offspring in its lifetime, after one generation there will be 2x200=400 worms on the plate and after two generations there will be 400x200=80,000 worms on the plate (density-dependence notwithstanding); after three generations there will be 80,000x200=16 million. There were many more than 400 and certainly fewer than 80,000, so we assume two generations is probably close to the true number of generations.

Assuming that *t* = 2, we find 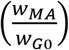 = 0.728, or in other words, relative competitive fitness declined by about 27% over 250 generations of MA. Scaled relative to the G0 ancestor, ΔM_*W*_, ≈ 1.09 x 10^−3^/generation. If *t* is closer to 3, 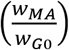 ≈ 0.81 and ΔM*_w_* ≈ 0.76 x 10^−3^/generation.

Averaged over the two blocks, the REML estimate of the among-line variance in log(*CI_H_*) increased from zero in the G0 ancestor to 0.534 in the MA lines, giving VM = 1.09 x 10^- 3^/generation (Table 2; Likelihood Ratio Chi-square = 23.7, df=2, P<0.0001). Scaled as a fraction of VE, the mutational heritability 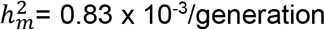. By way of comparison, the point-estimate of 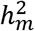 for non-competitive fitness in these lines under the same conditions is ~1.29 × 10^−3^/generation [30].

## Discussion

A simple but important finding is the close quantitative agreement between our estimate of the Zero Male Fitness mutation rate and that gleaned from the results of a previous MA experiment on the N2 strain [41]. Those estimates are subject to several sources of uncertainty, both experimental (e.g., perhaps if we had tried harder, we could have gotten fertile males from the lines that did not produce them) and biological (the distribution of mutational effects). The experimental uncertainty in this case leads to an overestimate of the ZMF rate. By way of comparison, the lethal recessive mutation rate in N2 has been estimated, from very limited data, to be on the order of ~0.01/generation [42, 43]. If we assume that each genetic death (lethal mutation) or male dysfunction (ZMF) is due to one and only one mutation, and that the estimate of 80 mutations per MA genome is not far off, then the fraction of ZMF mutations is 7/(80*60), about 0.15%. In every 100 genomes there will be ~32 new mutations, of which about one will be lethal, so the fraction of mutations that are lethal is 1/32, or about 3%. Thus, we infer that the fraction of ZMF mutations is around 5% of the lethal fraction.

What of the idea that male fitness is more susceptible to the cumulative effects of mutation than hermaphrodite fitness? Generally speaking, fitness is a function of survival, fecundity, and timing of reproduction. In the hermaphrodite assay, there was no discernible effect of mutation accumulation on the probability of reproducing (a finding which recapitulates the result from a previous non-competitive assay), so the decline in relative fitness with MA is encompassed by the ~0.11% per generation decline in relative competitive fitness. In males, both the probability of siring offspring and the fraction of offspring sired given that the male did mate successfully declined. A calculation based on the point estimates of the ΔMs (Table 1) shows that, after one generation of MA, hermaphrodite relative fitness will have declined by 1 - [(1 – 1.95×10^−5^)(1 – 1.09×10^−3^)] ≈ 0.11%. By the same reasoning, male relative fitness will have declined by 1 - [(1 – 6.82×10^−4^)(1 – 1.04×10^−3^) ≈ 0.17%. Thus, male fitness is certainly no less sensitive, and perhaps slightly more sensitive to the cumulative deleterious effects of mutation than is hermaphrodite fitness. This result is quantitatively nearly identical to the finding that male *Drosophila melanogaster* decline in fitness ~1.5X faster from mutation accumulation than do females [20].

The cumulative effects of selection depend on both the effects of an allele on fitness (the selection coefficient, s) and the effective population size, *N_e_.* Based on whole-genome sequencing of a subset of the MA lines in this experiment [44], the per-genome mutation rate is not less than 0.2 (one mutation every five generations) and unlikely to be more than about one mutation per generation, so the average MA line carries somewhere between 50 and 250 mutations; we think the true average is likely to be about 80 (AS and CFB, unpublished results). The cumulative decline in hermaphrodite relative fitness is about 27%. Thus, we can bracket the (arithmetic) mean homozygous effect on relative competitive fitness of new mutations, *s̅,* as lying somewhere between tΔM/50 and tΔM/250, where *t* is the number of generations of MA.

For hermaphrodite competitive fitness, tΔM_*w*_ ≈ 0.27, so the average selective effect *s̅* is bracketed between about 0.27/250 ≈ 0.1% and 0.27/50 ≈ 0.6%; if our estimate of 80 new mutations is correct, *s* ≈ 0.33%. For male relative fitness, tΔM ≈ 0.35, so *s̅* is bracketed between about 0.14% and 0.7%, with a best-estimate value *s̅* ≈ 0.44%.

*N_e_* of *C. elegans* has been estimated from the standing nucleotide diversity as being on the order of 10^4^ [10]. If a mutant allele is under selection in males but is neutral in hermaphrodites and males represent 1% of the population, the average selection coefficient on a mutant autosomal allele would be (0.99)(0) + (0.01)(0.0044) = 4.4×10^−4^, on the cusp of effective neutrality. Deleterious alleles with selection coefficients *s* ≈ 1/*N_e_* are the most pernicious with respect to population mean fitness [45]. On the face of it, it would appear that males in *C. elegans* are in peril of mutating their way out of existence. However, that conclusion is based on the strong assumption that mutations that affect male fitness have no pleiotropic effects on hermaphrodite fitness.

This study has two limitations. First, we would like to have an estimate of the mutational correlation between male fitness and hermaphrodite fitness, because those data would illuminate the extent to which intersex pleiotropy (“intralocus conflict” if effects are antagonistic between the sexes, [46]) is an inherent feature of genomic architecture, without the confounding influence of natural selection. However, the lack of significant mutational variance in either of the male fitness traits (*П_μ_* and *CI_M_*) obviously means the estimate of any covariance with those traits is zero. The male-fitness assay included fewer MA lines (53) than did the full hermaphrodite assay (80), although each block of the hermaphrodite competitive fitness assay included fewer lines (~40 vs ~50) and the estimates of VM were highly significant in each block (LRT, P<0.001). We have assayed many hermaphrodite traits with ~50 250-generation MA lines and detected significant VM (e.g., [14, 47]). Mating is inherently subject to many more sources of variation than is internal self-fertilization, and the results reflect that greater variability.

The second limitation is that males compete for fitness not only with other males, but also with the hermaphrodite itself. Measuring male-hermaphrodite competitive fitness in our context requires a recessive marker in the hermaphrodite competitor, so that the offspring of a cross can be distinguished from the hermaphrodite’s self-progeny. Unfortunately, we were unable to find a recessive marker that had reasonably high fitness and could also be reliably scored at sufficiently high throughput to enable a full assay, either with the worm sorter, by image analysis, or by eye.

Male-male competitive index includes both a behavioral component and a sperm-competition component, which cannot be discriminated with our assay. Both features could potentially affect male-hermaphrodite competitive fitness. A hermaphrodite paired with a male that is a poor mater may sire a larger fraction of offspring prior to exhaustion of its sperm than will a hermaphrodite paired with a good mater. Similarly, a hermaphrodite mated to a male with poor sperm will presumably sire a larger fraction of offspring than a hermaphrodite mated to a male with good sperm. Male sperm generally outcompete hermaphrodite sperm [48], so if it is assumed that the entire difference in male-male competitive fitness is due to reduction in sperm-competitive ability and that wild-type hermaphrodite sperm would be no better competitors than wild-type male sperm (which seems reasonable), then the strength of selection against male-male competitive fitness provides an upper bound on the strength of selection acting on the competitive ability of male sperm relative to hermaphrodite sperm. Alas, no such simple approximation is possible with respect to male mating behavior, because the fitness consequences of even the simplest aspect of male behavior, time to mating, depend on the distribution of timing of hermaphrodite self-fertilization.

To conclude, the results of this study indicate that selection acting on mutations affecting male function is similar to, or perhaps slightly stronger than, selection on mutations affecting hermaphrodite function. However, a full accounting of mutations affecting the full spectrum of components of male fitness remains incomplete.

## Acknowledgments

We thank Dustin Blanton, Whitney Bour, Myrnelle Damas, Thomas Keller, Laura Levy, Gigi Ostrow and Naomi Phillips for assistance in the lab. *C. elegans* strains were graciously provided by Keith Choe (UF) and the Caenorhabditis Genetics Center. Support was provided by NIH grants R01GM107227 to CFB, E. C. Andersen and J. M. Ponciano, and S1010OD012006 to CFB, L. Bianchi, K. Choe, and A. S. Edison.

---

**Supplementary Table S1a.**
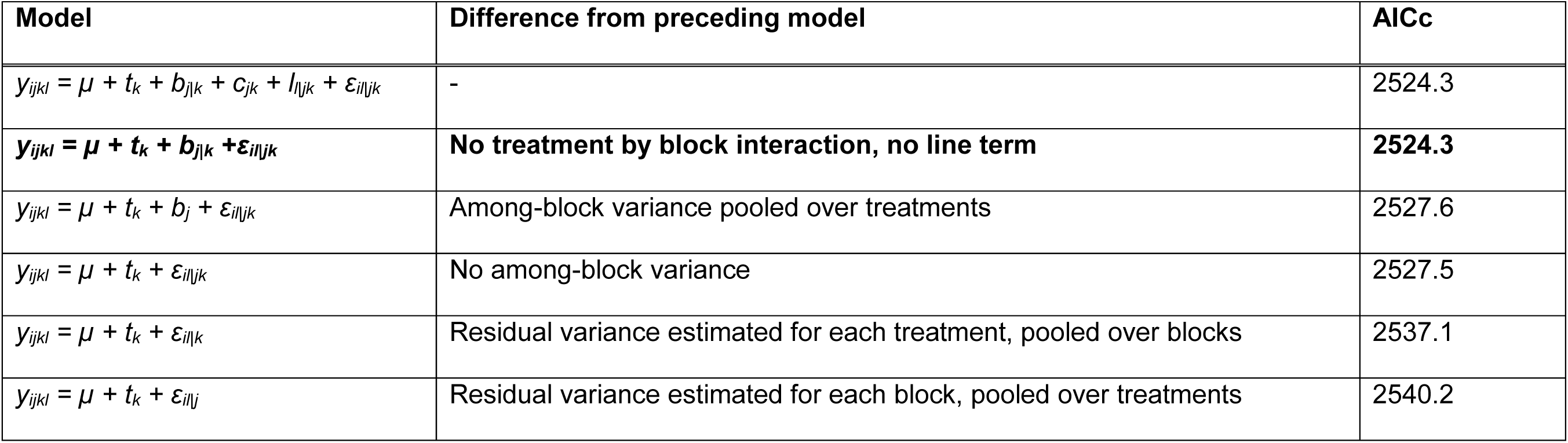
AICc scores of linear models tested for male CI (*CI_M_*). *μ* is the overall mean, *t_k_* is the fixed effect of treatment *k* (G0 or MA), *b_j_* is the fixed effect of block *j, c_jk_* is the fixed effect of the treatment by block interaction, *I_l_*_\*jk*_ is the random effect of line (or pseudoline) *l*, conditioned on block and treatment, and *ε_l_*_\*jk*_ is the random residual, conditioned on block and treatment. Terms with a REML estimate of zero were removed from the full model prior to testing simpler models. The best-fit model is shown in bold type.

**Supplementary Table S1b.**
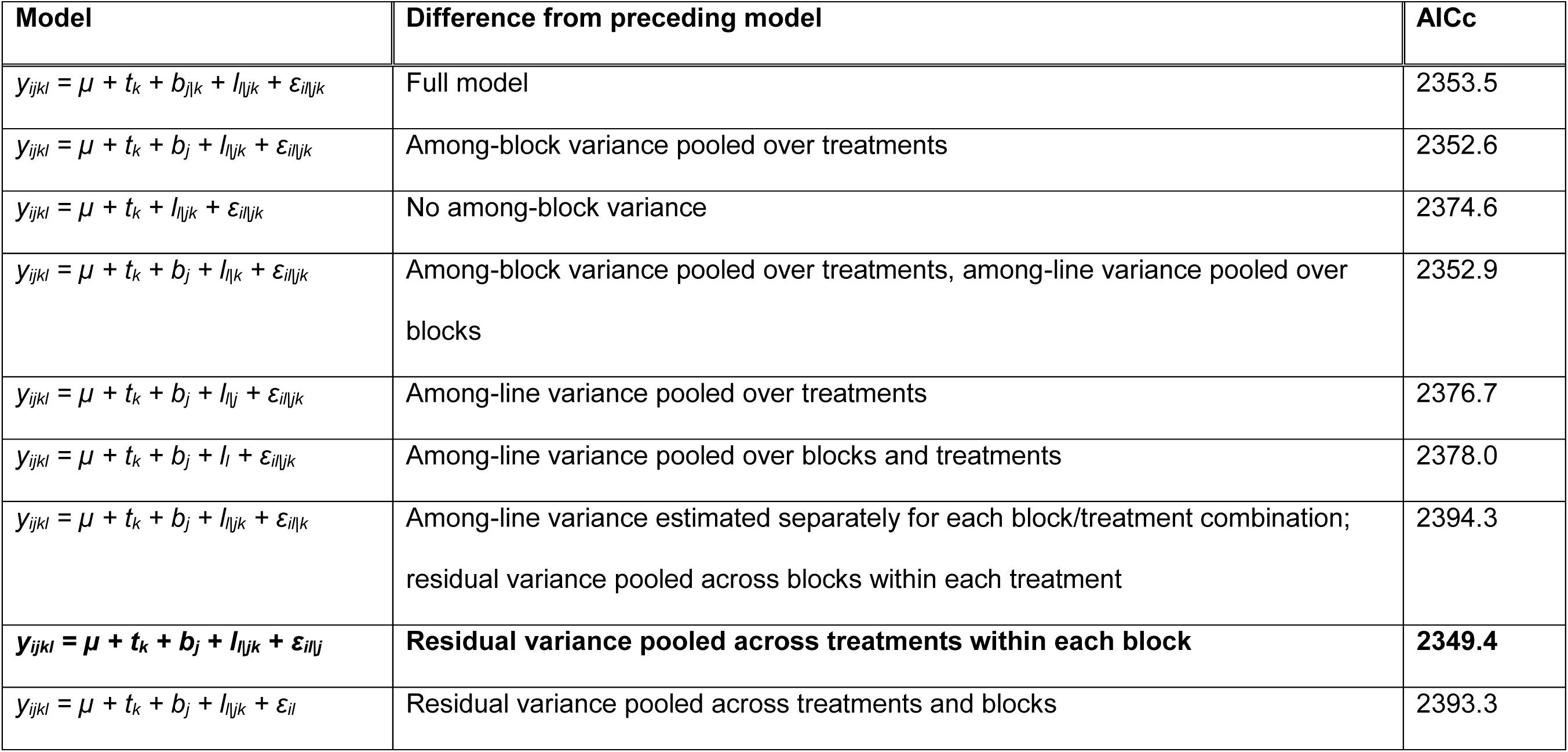
AICc scores of linear models tested for hermaphrodite CI (*CI_H_*). Definitions of variables are the same as in Table S1a. Note that since each MA line was present in only one block, there is no treatment by block interaction in these models. The best-fit model is shown in bold type.

**Supplementary Figure S1.**
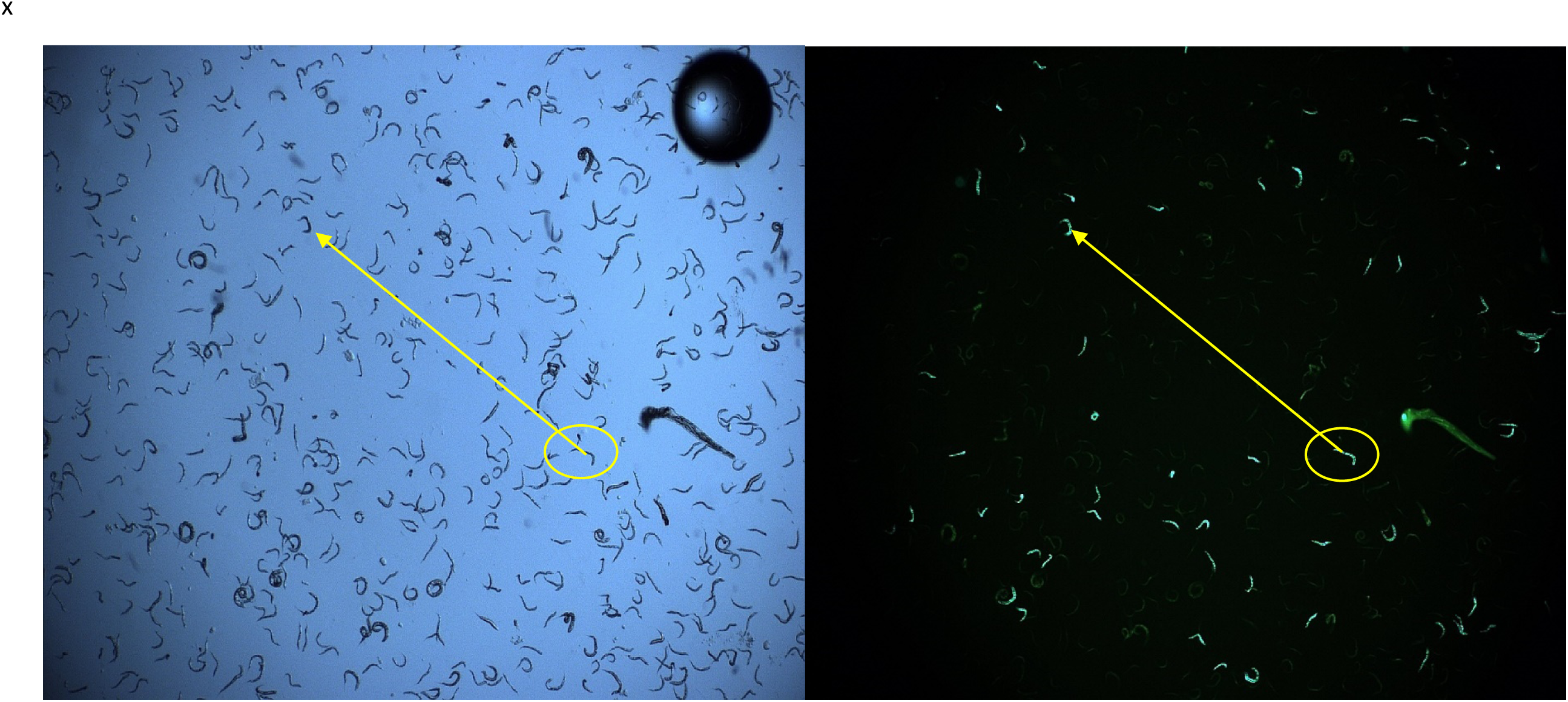
Example of a side-by-side comparison of the same sample of worms taken under transmitted white light (left) and under 470 nm fluorescent light (right) in the hermaphrodite competitive fitness assay. Worms that appear green under fluorescent light are the GFP-marked competitor (ST-2); worms that do not fluoresce are the focal type, either MA or G0. The yellow oval and arrow highlight the same individual worms in the two images and are shown to emphasize that the images are an exact overlay of each other. The difference between the number of worms in the left image and the right image is the fraction *p* of the focal type.

**Supplementary Figure S2.**
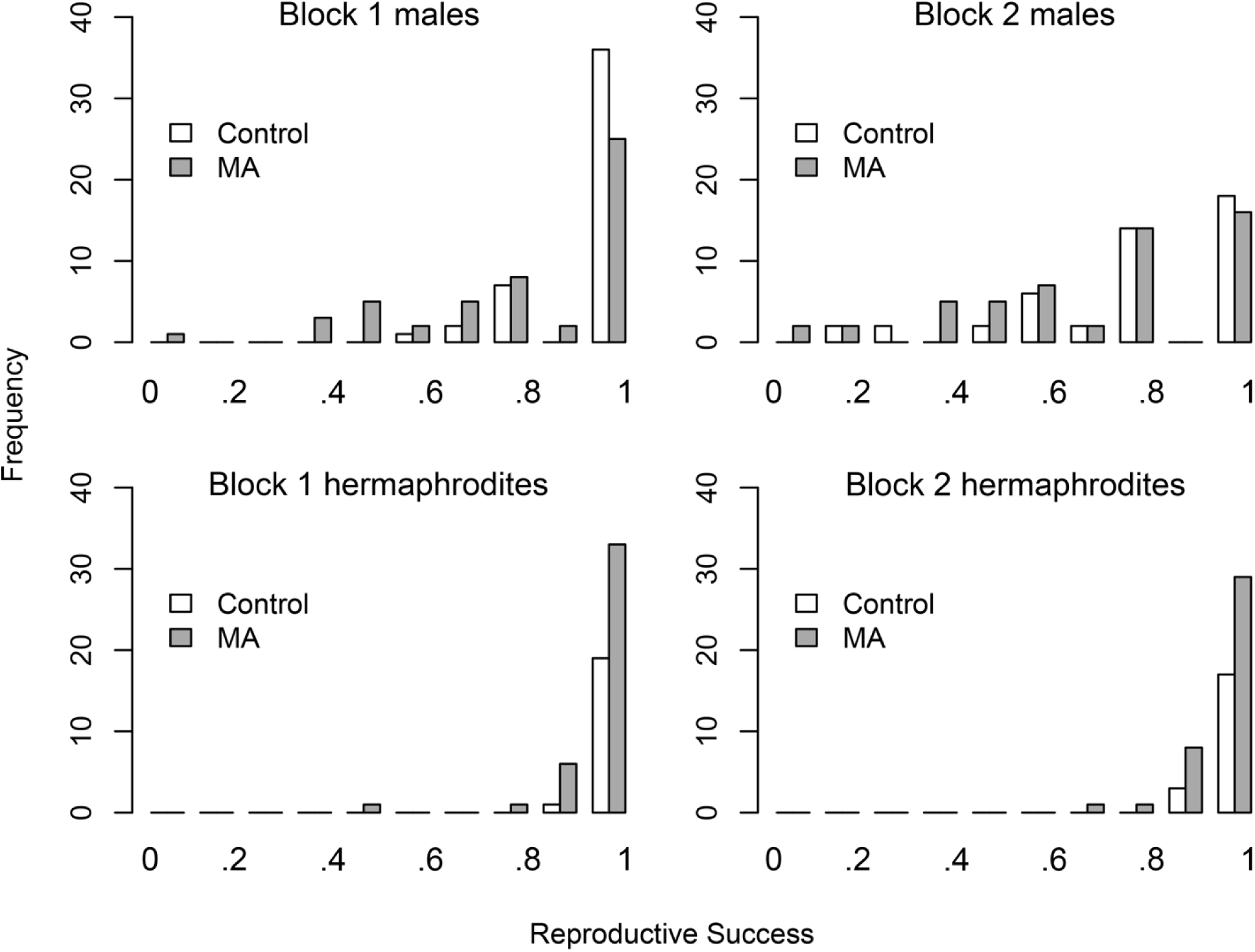
Distributions of line means of reproductive success (= mating success in males); X-axis values are probabilities of reproducing. Upper panels, male reproductive success (*π_M_*); lower panels, hermaphrodite reproductive success (*П_Η_*).

**Supplementary Figure S3.**
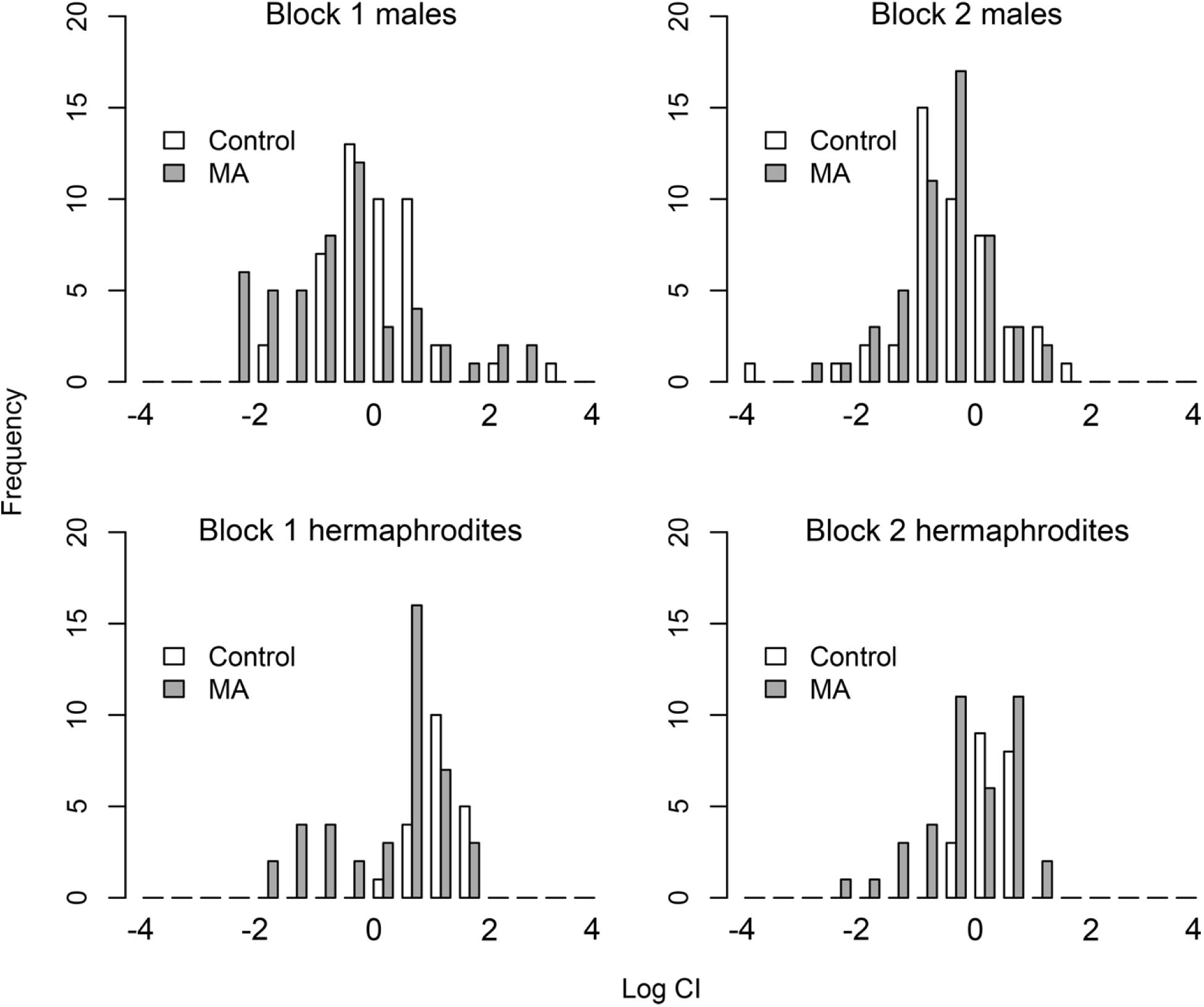
Distributions of line means of log(*CI*). See Methods for description of calculation of *CI.* Upper panels, male reproductive success (*CI_M_*); lower panels, hermaphrodite reproductive success (*CI_H_*).

## Supplementary Appendix A1

Counting worms with the BioSorter™

The Union Biometrica BioSorter™ is a flow cytometer equipped with a flow cell of diameter large enough (250 μm) to permit worm-sized particles to be detected; the specifications are included in the manufacturer’s documentation at http://www.unionbio.com/biosorter/. The Biosorter registers an event (a worm, piece of debris, etc.) when a laser detector detects a reduction in optical density (“extinction”) relative to the background optical density. The extinction profile (the time of flight and area under the curve) is correlated with the length and shape of the object passing the detector, enabling worms to be distinguished from debris and other non-worm objects. The Biosorter is equipped with a 488 nm excitation laser and can detect GFP fluorescence. Events can be “gated” according to time of flight, extinction, and intensity of fluorescence. Worms at the L2 stage of development or larger can be distinguished from non-worm events such as debris with reasonably high confidence (details follow). Eggs and L1 stage worms have a much lower signal to noise ratio, so we exclude those developmental stages from the analysis.

Worms were washed from the competition plates (see Methods in the main text) in approximately 1.5 ml of M9 buffer into 1.5 ml microcentrifuge tubes. Tubes were centrifuged at ~2K x g for one minute, the supernatant decanted, and the pelleted worms resuspended in 100 μl of M9 and pipetted into a well in a 96-well microtiter plate to be counted with the Biosorter.

The contents of each well of the 96-well plate was counted using the LP Sampler™ microtiter plate sampler.

The sample may contain bacterial clumps, shed worm cuticles, and other non-worm debris that can register as false positive events in a “worm” gate. Because these false positives are not fluorescent (beyond a certain background level), and we count both fluorescent and non-fluorescent worms, false positives inflate the apparent fraction of non-fluorescent worms in a sample. Overestimation of the fraction of non-fluorescent worms, *p*, leads to an overestimate of the competitive index *p*/(1-*p*). To quantify and correct for false positives, we set up 125 35 mm NGMA test plates with five different population sizes (50, 100, 150, 200, 250) at five fluorescent-to-non-fluorescent-worm ratios (1:4, 2:3, 1:1, 3:2, 4:1), each replicated five-fold. Worms were sorted from a mixed-stage mass culture onto the test plates using the BioSorter in “sort” mode using the same gates as in the fitness assay. Worms were washed from the plates into a well of a 96-well microtiter plate and counted as described above.

The sorting efficiency and rate of detection of fluorescent worms are both > 99%. Some worms are lost in the wash step, and not all worms present in a microtiter plate will be recorded as events based on extinction (as opposed to fluorescence). However, there is no reason to think that loss or failure to record are genotype-specific. The false-positive rate is calculated as follows. Terms in bold text are depicted in Supplementary Figure S4 below.

> **N_T_ = Total events recorded in the test sample**
>
> N_W_ = Total events recorded in the **worm gate**, gating on extinction.
>
> N_FS_ = Number of fluorescent worms sorted onto the test plate
>
> N_FR_ = Total fluorescent events recorded in the worm gate, gating on fluorescence.
>
> N_W_-N_FR_ = [number of wild-type worms + non-worms in the worm gate]
>
> L = [N_FS_ - N_FR_]/N_FS_, Loss Rate, i.e., fraction of worms lost in washing and counting in LP Sampler
>
> R = 1-L, Recovery rate
>
> FP = [(N_W_-N_FR_)-N_FS_XR]/(N_T_-N_FR_) is the estimated false-positive rate.
>
> These calculations were applied to 96 of the test plates, leading to an overall estimated false-positive rate of 9% (FP=0.09). The number of non-fluorescent worms is estimated as number of non-fluorescent events in the worm gate,(N_W_-N_FR_)(1-FP).
>
> In assay block 1, worms were stored at 4° C for four days prior to counting. To account for the potential effect of refrigeration and storage, 29 of the 125 test plates were stored at 4° C for four days prior to counting. On average, the number of fluorescent events recorded after four days of cold storage (N_FR,C_) declined, leading to an increase in the loss rate after cold storage (LC). We assume the difference between L and L_C_ (and thus R and RC) is due to dead worms that no longer express GFP but that still register as events when gated on extinction. In block 1, the number of fluorescent worms in a sample was estimated as N_FR,C_+N_FR,C_X[(R-R_C_)/R_C_].

**Supplementary Figure S4.**
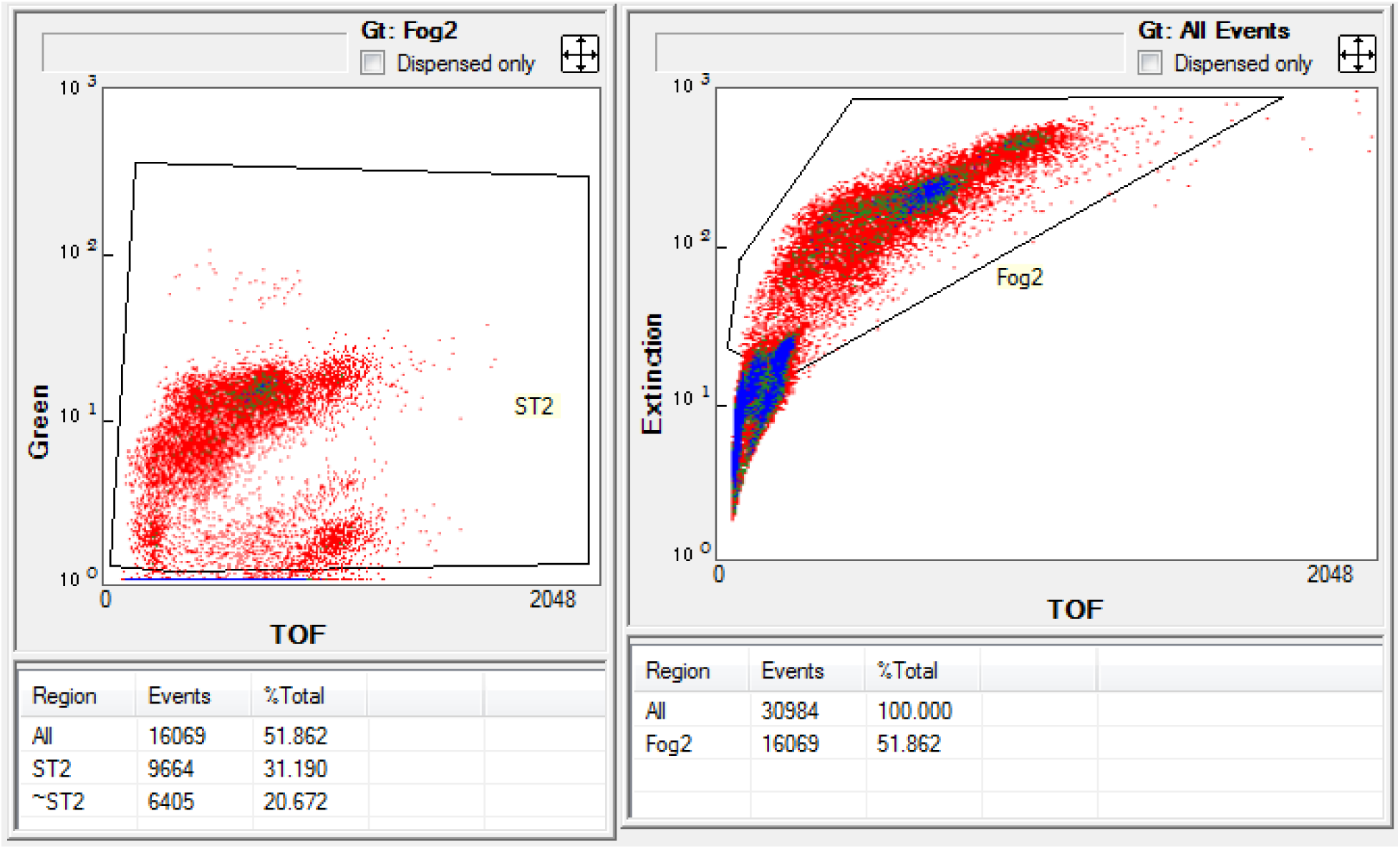
The plot on the right displays a density plot of all the events recorded in one well of a 96-well plate, with log(extinction) on the y-axis plotted against time of flight on the x-axis (TOF is a proxy for object length). The density of events increases from red to green to blue. The polygon enclosed by the black lines is the “worm gate”, designated “Fog2” in this figure. Events characterized by extinction/TOF ratios within the gate are classified as worms; events falling outside the gate are classified as not worms. In this example, the total number of events recorded in this plate is 30,984, 51.86% of which fell in our worm gate. The plot on the left is a subset of the worm gate (designated “ST2” in this example) and shows a plot of log(intensity of green fluorescence) against TOF. In this example, 9664/16069 events are classified as fluorescent (and thus as worms of the ST-2 strain).

